# Cyperus rotundus root extract inhibits progress of lymphedema in mouse tail model

**DOI:** 10.1101/2021.10.15.464519

**Authors:** Nikhil Pandey, Priyanka Mishra, Yamini.B. Tripathi

## Abstract

Lymphedema is clinically manifested as swelling due to abnormal accumulation of interstitial fluid attributed to inefficient fluid uptake and reduced lymphatic flow. Here we have evaluated the effect of Cyperus rotundus root (CRR) ethanolic extract in a mouse tail model of lymphedema and hypothesized that *Cyperus rotundus* plant known for its anti-inflammatory effect through inhibition of TNF alpha will be effective in managing this condition. The skin was removed after leaving 1cm of distance from the base of the trunk. Cut was introduced in sterile condition. The animals were divided into Experimental control(EC) and Cyperus rotundus (CRR) treated groups. A change in tail volume around the wound was monitored upto the 20 days. Kinetics of the swelling was calculated for statistical significance. Further TS of upper part of the wound was stained with H&E stain and documented for histological changes

**RESULTS:** In EC group, gradual rise in swelling was recorded, which peaked on 10^th^ day and continued up to 15^th^ day, followed by gradual decrease. In CR extract treated group, the swelling was significantly low and peak was obtained on 8^th^ day, while in EC group the swelling was significantly higher and peak was obtained 11^th^ day. The histological section show, fibrous band intersection the lobules of adipocyte and lymphatic vascular channel and muscles in the sub epithelium region in the EC group, which was very less in CRR group.

**CONCLUSIONS:** Polar fraction of CRR significantly prevents the progress of lymphedema and which lays an important role of this plant as for the drug development in future.

## Introduction

The lymphedema is a pathophysiologic progression characterized by the buildup of protein-rich fluid or lymph fluid in the interstitial space, local adipose deposition, and tissue fibrosis causing from the non-congenital factors such as injury, infection, obstruction, or congenital defects in the lymphatic system. It is explained that all edema in the extremities is created from a relatively incompetent lymphatic system that is overwhelmed by fluid microfiltration in the tissues.(1) This can either be from an inadequate lymphatic system that cannot bear the load of a normal amount of fluid in the extremities (lymphedema) or from a normal lymphatic system that is overwhelmed with a high microcirculation fluid accumulation from a myriad of possible conditions, such as heart failure, kidney failure, liver disease, and malnutrition to name a few (2).Cancer and its treatment is another reason for the development of the secondary lymphedema due to blockage of lymph node in the presence of tumour or removal of tumour along with the lymph node cause to fluid to build-up at near tissues.(3) Numerous signalling pathway are involved in pro-lymphogenic effect. Major studies has been concentrated on VEGF-c/VEGF -3 pathway. Regulation, proliferation and migration of the lymphatic endothelial cell is done by VEGF -C (4)(5).Its overexpression results in progression of the lymphogenesis and hence it is exploited as ideal therapeutic taget for the progression of secondary lymphedema. Another growth factor like VEGF-A shows is its pro-lymphogenic effect and regulates remodelling of the of the lymphatic vessel(6).Apart from this, TNF alpha(7)(8) and TLR signalling pathway (9),COX-2 as well as prostaglandin E2 receptor signalling(10) are involved in the progression of the lymphedema. To encounter it, presence of anti-lymphangiogenic effect like cytokine like(IFN-γ) which are produced by numerous cells such as the T-cell and in some extent by macrophages have the effect of acting against the pro-lymphangiogenic by modulating the Janus kinase signal activating the JAK STAT transcription pathway. Besides the T-helper (Th2) cytokines such as IL-4,13(11),and TGF-β1 (transforming growth factor beta 1) shown to be anti-lymphangiogenic in nature(12,13,14)

One of the inflammatory mediator which is an increased Leukotriene B4,in the lymphoedematous tissues also has substantial anti-lymphangiogenic effect but its effect are dependent on the concentration in the low concentration it promotes the lymphangiogenesis and at high concentration its role get reversed and act as anti-lymphangiogenic (15).

Balance between the pro and anti-mechanism of lymphedema is maintained by the these cells and cytokines. Pro-lymphangiogenic molecules like VEGF-C, VEGF-A, B-lymphocytes(16) (17), Fibroblast type reticular stromal cells(18) and CD11b macrophages(19) (20)(21). Similarly anti -lymphangiogenic effect are produced by the th1 and th2 cytokines of the T cells(22). Hence a varity of cells can initiate the diverse signalling pathway which have the important effect on the endothelial cell of lymphatic vessels and its balance can regulate or impair the regeneration of lymphatic vessel (23) (24) Henceforth, the mechanism that can utilised these will be the potential targets for the use of therapeutic intervention.

Currently in the developed world, a major number of cases of the lymphedema are linked to secondary lymphedema and it is caused by the blockage of the lymph flow by malignant tumor or due to presence of the injury like a fracture or injury in the surrounding tissue of the lymphatic vessel. Similarly, the filariasis which is one of the widespread parasitic diseases, causes the obstruction of the lymphatic system resulting in the development of enlargement of legs (elephantiasis) (25), chronic wounds and in arms, scrotum and breasts like human, other affecting filariasis are onchocerciasis, loiasis and mansonellosis. (26)

Being the progressive disease its early diagnosis and treatment is vital. Currently the treatments are present in the form of post -management of lymphedema such as Decongestive lymphedema therapy (DLT),Manual lymph drainage (MLD) and compression, as per the drug therapy, which are the adjunctive for the pain or secondary infection are present(27). One of the biggest challenges in managing and monitoring lymphedema is volume measurement. Currently, there is no standard directions for diagnosing and monitoring the progression of lymphedema. Circumference measurement is the most frequently used technique due to the low expense and ease of use. Circumferential measurements are either taken at bony landmarks or established locations along the limb(28).

Hence at present the use of the pharmacotherapy agents in clinical and experimental studies are focusing on inhibiting inflammation like inhibiting cyclooxygenase leading to reduce tissue inflammation, such anti-inflammatory compounds(29). Example like use of selenium (Sodium selenite) is has shown to reduce lymphedema volume (30)but this also bring the adverse effect of it such as gastrointestinal symptoms (e.g. Nausea, vomiting and diarrhoea and irritation of the respiratory system. Since several etiogenesis of lymphedema are reported in literature which is

1. High oxidative stress causing the persistent enhanced formation of reactive oxygen species (ROS) and leads to the upregulation of lipid peroxidation in chronic lymphoedematous tissue. The strengthening of antioxidative defence mechanisms could be useful in the therapy of chronic lymphoedema. Due to high ROS environment causes the expression of pro -inflammatory gene upregulation in lymphedema patients and in animal model such as CD 14,IFN-γ (interferon-gamma) receptor, tumour necrosis factor-alpha (TNF-α), integrin alpha 4 beta 1 (α4β1; also known as Very Late Antigen-4 or VLA-4), tumour necrosis factor receptor p55 (TNFR1), and CD44.(31)
2. The secondary Lymphedema which is more common (32) and prevalent shows the involvement of several cofactors in the development such as congestive heart failure or renal diseases. This has been further suggested that various genes like VEGFR2, VEGFR3,RORC,GJC2 and FOXC2 are possibly involved in the developing of secondary lymphedema following the breast cancer therapy. (33)
3. Since the present line of treatments includes non-invasive and invasive methods for lymphedema such as therapy (DLT,MLD, Compression) and Microsurgical techniques like vascularized lymph node transfer (VLNT),Suction-associated protein lipectomy (SAPL) in patients(34) Currently the recent improvements studying the pharmacotherapy and surgical treatments have revealed some promise.(35)(36)(37)
4. Currently in the, management of secondary lymphedema focus on the post -management of it via therapy or surgical intervention, but focus should be on the prevention and this could be done with the exploration of the new therapeutic drugs. These pharmacological intervention on the different pathways of inflammation, adipose tissue and change in the permeability of vessels could slow down or prevent the progression of the secondary lymphedema Therefore the finding of new natural drug from the plant, we started with the extract of Cyperus rotundus which is already in clinical use under the traditional Indian ayurvedic system for its preventive action of the reduction of obesity and the reducing of the blockage in vascular system. (38)
5. Therefore the treatment for this disease has yet to be characterized and the ideal treatment for this disabling disease has yet to be identified and likely includes a multimodal approach. One of the approach is animal models.

## Mouse Model

Many of the current knowledge in in the domain of lymphedema is fostered upon the earlier work of Olszewski (39) who developed a canine based lymphedema model in his experimental studies. Till now many animal based model has been utilised ranging from rodents to larger animals such as sheep and pigs in an attempt to closely study the human disease (40)

A consistent and reproducible lymphedema animal model is quite essential for such studies. Hence the procedure for microsurgical removal of the lymphatic vessels from the tail of the mouse causes the process of stagnation of the lymph and dilation of lymph vessel with the notable increase in the tail volume. Following the Collection of fibroblasts, fat, and skin cells, (41) weakened clearance of immune cells from the tail, and profound accumulation of inflammatory cells. These hallmarks features the acquired lymphedema in humans (42) For the mouse tail model, the blocking of the lymphatic collateral vessels present at both sides of a rodent tail is utilized to understand the underlying mechanisms of lymphedema induction is the recent most method for the studies related to lymphedema.(43). This model has the advantages of low cost and easy in handling, and its efficiency in interpreting the molecular mechanisms of lymphedema; however, as the pattern of lymphatic flow and regeneration in rodent tail is different from that of humans, there is a necessity of an animal model that mimics the secondary lymphedema of humans. Since this tail model closely simulate the histologic features of post-surgical lymphedema in humans (rather than primary or infection -related lymphedema) hence it is useful for the study of the inflammatory responses incited by the lymphatic injury therefore important for the understanding of the lymphedema pathophysiology.

### Cyperus rotundus

Cyperus rotundus, (Cyperaceae), known as Nagarmotha in India and Nutgrass as in common name. CR is known for its pharmacological properties like anti-oxidant.anti-inflammatory, anti-pyretic, wound healing properties. (44) (45).

It also reduces the lipid peroxidation, as measured by reduced thiobarbituric acid–reactive substances (TBARS) in several tissues including in rat-brain-mitochondria, induced by FeSO4 (46). The high flavonoids and polyphenols contents in the polar and non-polar extracts of Nutgrass are attributed to this property. For the first time we are reporting the effect of Cyperus rotundus on the mice tail lymphedema which has shown the promising result. Therefore, in future the more evidence will be beneficial for the conclusion of its potential for the use of CR in lymphedema management and the use of herbal remedies for lymphedema management. As in stage, from herbal extract (where due to presence of multiple compounds in crude extract it can be largely beneficial depending upon the dose or amount to be administered if properly regulated better efficacy can be shown,) to plant synthesised secondary metabolites stage is necessary for screening of therapeutic drug from these plant secondary metabolites (47). Hence Cyperus rotundus roots extract and its efficacy will provide helpful resource for more screening of its secondary metabolite to get more specific answers for the cure of lymphedema.

## Material and Methods

1. **Animal Procurement**- Female mice (Swiss albino) obtained from the Central Animal facility of Institute of medical sciences, Banaras Hindu University, Varanasi, were maintained at normal room temperature with 45%-55% of the humidity. The normal diet was provided during the course of the whole experiment.
2. **Preparation and standardization of the extract-**Cyperus rotundus roots (CRR) were procured from ayurvedic pharmacy of the institute and its authenticity was verified on standard pharmacogenetic parameters, according to the Indian pharmacopoeia The roots were powdered and then exhaustively extracted with Ethanol in continuous Soxhlet extractor for 36hrs. The yield was 17.5 percent. The CRR extract was given in the dose of 80mg.kg.BW/Per day. It was given orally with the help of gavage tube to the animals of the lymphedema group up to the time period of day 5, day 10, day 15, day 20. The experimental control group were given 1ml of drug vehicle (20% of Tween 80 in the water).
3. **Cyperus rotundus extract analysis (HRMS Analysis) -**In the analysis of CR extract, it was used to performed by utilizing QTOF instrument therefore we analysis we have found the 58 compounds present in the methanolic extract (Table 5)
4. **Tail lymphedema eexperimental design –** We carried the study for an acute post-surgical lymphedema in the tails of swiss albino mice female in three groups (i.e., Group I was Experimental control, and Group II for Cyperus rotundus (80mg/kg/BW). The swiss albino mice /females were chosen for this study of similar weight.(20-25gm) The skin was removed was done after leaving 1cm of distance from the base of the trunk. Cut was introduced in sterile condition. The administration of ether was applied for the anaesthesia and skin and subcutaneous tissue between from the base of 5-10mm to the distal region of the tail was removed (fig1 & 2). The mice were kept in separate cages to avoid the damage being caused by the other mice. We measured the volume of the tails every day for 20 days, thickness and histological studies were also done to assess the lymphatic adequacy.The tail was ethically amputated in aseptic condition on due time period for the examination of tail for mRNA expression and Histological studies for its gross examination of the tail. During the given time scale the photography of the tail was carried out to examine the change in inflammation in lower and upper part of the cut. The measurement of tail was done with digital caliper. This study was done after receiving the animal ethical approval from Institute of medical sciences, BHU, Varanasi.
5. **Volume measurement** –At post - surgery every mice were taken for the measurement of tail diameter of by digital vernacular calliper of upper and lower ends of the swelling. Along with the a ruler for measuring the length, digital vernacular calliper was used to find the thickness increase and decrease in the murine tail. Photos were taken to measure the diameter of each region. Besides after converting the diameter into circumference

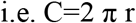

the next formula used to convert it into volume by regarding as sliced cone (48).

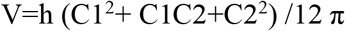
6. **Histological assessment** – The collected tail samples were fixed in formaldehyde and was dehydrated, embedded in paraffin, sectioned into slices (5 μm) and stained with haematoxylin and eosin (H&E).
7. **Statistical analysis -** Graph pad (8.0.2) was used for the statistical analysis of mouse tail volume and circumference. The measurement of the volume change was calculated alongside with their mean and standard deviation in both the groups. The degree of volume changes s measured between two groups. Results are expressed as Means and SD with the p-value<0.001 in volume, upper and lower circumference of both the groups. Statistical significance was analysed with unpaired student’s T-test for comparison between the 2 groups.

## Results

1. **Phytochemical analysis**- The LCMS data reveal the presence of the 58 compounds which are significant for its medicinal
2. Effect of Cyperus rotundus on the tail lymphedema. **Change in volume**-The gradual increasement of the tail circumference and total volume at post-surgical was observed After the assessment of the diameter and volume of the mice’s tail in both CRR and EC group the CRR group shown the significant decrease in volume compared to volume of the experimental group. Significant difference was observed and its continuation in the volume of the experimental group and Cyperus group from day 6 after the tissue incision (P<0.01), and this difference was continued till the end point of the experiment. When observing the volume change, the most distinct increase was observed in day 11 where highest difference was observed when comparing the Exp.Control to CRR group. The CRR group shown to attain the highest swelling area on the day 8 and day 9 whereas the Exp.Control shown the highest swelling on the day 11^th^. Continuous decrease in volume was presented after day 9 for CRR and day 11 for Exp.Control. Furthermore, according to the assessment of the diameter the higher swelling was observed on the upper (Proximal) compared to lower (Distal) area in all the mice of the two groups.(Table No.4) **Histology Studies-** The gradual increasement of the tail circumference and total volume at post-surgical was observed. Histological sections showed the inflammatory response and lymphatic microvascular dilation. The CRR group (n=10) has shown the swelling of the tail in its highest form at day 8 and day 9, in contrast to Experimental group (n=10) shown the maximum swelling at day 11 and Day 12.

## OBSERVATION

1. We carried out.an experimental model of the post-acute-surgical based lymphedema. In the tails of female Swiss albino mice. Removal of skin and subcutaneous tissue in tails of the mice (10mm or 1cm from the base of the tail). Following this, the murine shown the acquired lymphatic swelling. We measured the volume on every day for 3 weeks to assess the lymphatic flow. We also observed during our studies that necrosis could be effectively avoided by paying attention to many details in the modelling process. The hair follicles on the inflamed area were erected and the shiny texture was started to appear as the marked area started to swell. Inflammation area of the tail near the edema were hairless and this condition was continuous in both the upper and lower area of the edema area.

## DISCUSSION

Since from the past to present many research has been conducted on lymphedema but the mechanism for histological changes from the cause to molecular mechanism remains still in debate. The most common treatment which is still used is a non-drug treatment, known as decongestive physiotherapy (49) which is a complex one, but this deals to the post management, not the cure. Since the core question lies in the pathogenesis of lymphedema and its progression. This is an important domain to identify and utilise new therapeutic targets as currently there is no drug available in common person domain. So far, the benzopyrones are the sole drug for the treatment of lymphedema, which act by removing the excess amount of protein via causing the proteolysis by macrophages, as result causing the reduction in the swelling, fibrosis and chronic inflammation. (50). Besides the other anti-inflammatory drugs are such as the topical application of tacrolimus which as an anti-T cell immunosuppressive drug is used for the prevention of lymphedema development (51). The Uncovering of the new compounds which can act on the various different pathways during the lymphedema is utmost important task for patients. Towards this objective here for the first time we have tested therapeutic response of CRR. But in future its specific role on the molecular target involved in etiopathogenetic of lymphoedema will be carried out. In this research experiment we have taken Cyperus rotundus root as natural candidate to create the domain for its more thorough work in the domain of lymphedema, and how its specific important metabolites imparting its role in decreasing of the lymphedema in the mice tail for the start of the new domain. In future its detailed molecules can show a specific pathway on which it is acting, to derive ideal candidate as natural compound. As CR shown to be potent medicinal plant, showing good efficiency as anti-oxidant and anti-inflammatory agent imparting its role via scavenging superoxide radicle, hydroxyl radicles due to presence of high flavonoids and polyphenols in polar and non-polar extract of CR.(Ref). Compounds like (+)-Nootkatone and (+)-valencene from the rhizome of the CR shown to enhance the rate of survival in septic induced mice because of the presence of Heme oxygenase -1 induction (52). To make it more robust medicinal plant, CR has shown vital action on the HO-1 by inducing it, and as result it HO-1 Provide cytoprotective, anti-apoptotic and immunomodulatory effects. CR role in inducing HO-1 also provide a beneficial role in the modulation of pro-inflammatory stimulators as inducible nitric oxide synthase (iNOS) Tumour necrosis factor-alpha (TNF-alpha)(53). Effect of Cyperus rotundus roots in the decreasing of the swelling in mice tail of our experiment can also be supported by the Earlier reports from the in-silico studies have indicated the role of CRR as inhibitor of 5-lypoxygenease and leukotriene A4 hydrolase (LTA4H) done by Fares Fenanir Leukotriene B4 (LTB4)(54) is one of the most important inflammatory lipid mediator in the pathogenesis of lymphedema. These are biologically active lipids, mainly produced by pro-inflammatory cells such as macrophages, eosinophils, mast cells, neutrophil, as result these lipid mediators generate the inflammatory response by binding and activating their G-protein coupled receptors (55) Hence in our work the result of the Cyperus rotundus in mice tail lymphedema provides another evidence. Apart from this, another feature of lymphedema which is the deposition of adipose tissue which may triggered by subtle dysfunction or the injury in the lymphatic system (Which is done polarization of infiltration M1 into adipose tissue makes the entre environment, inflammatory and attributing the blockage of the lymphatic system blockage. In this process T -cell activated to release the molecular pathway(56). Thus, inflammation and the deposition of fat are closely in linked in the lymphedema progression. In the same way, expression of IL-6 is upregulated in progression of lymphedema which is an important interleukin for the maintenance of homeostasis of adipose tissue and also in obesity pathophysiology (57). The lymph fluid contains higher IL-6 concentration compares to the fat from the subcutaneous area, which positively indicates the involvement of lymphatics in the clearance and transport of inflammatory cytokines(58).Hence this conclude the correlation of the fat deposition with the IL-6 expression in an inverse fashion and loss of this IL-6 connected to lymphatic injury making a sharp upregulation of fat depositions (59).

Cyperus rotundus root extract effect in the decrescent of swelling is strongly supporting its earlier described role as downregulating of pro-inflammatory cytokines genes. Molecule’s present in CRR has been reported to inhibit the pro-inflammation via the inhibition of the NF-κB and STAT3 pathways and ROS. (60)(61)

## Conclusions

Although research on lymphedema requires understanding through pathophysiological analysis, experimental research remained inactive due to lack of experimental subjects. We have a Swiss albino mice tail model of secondary lymphedema, which can be used to stimulate an trait of human lymphedema and provides the insights of functional and structural changes of lymphedema.

## Acknowledgement

**NP are thankful to ICMR for providing the financial assistance**.

## SUPPLEMENTARY DATA

**Fig.1.**
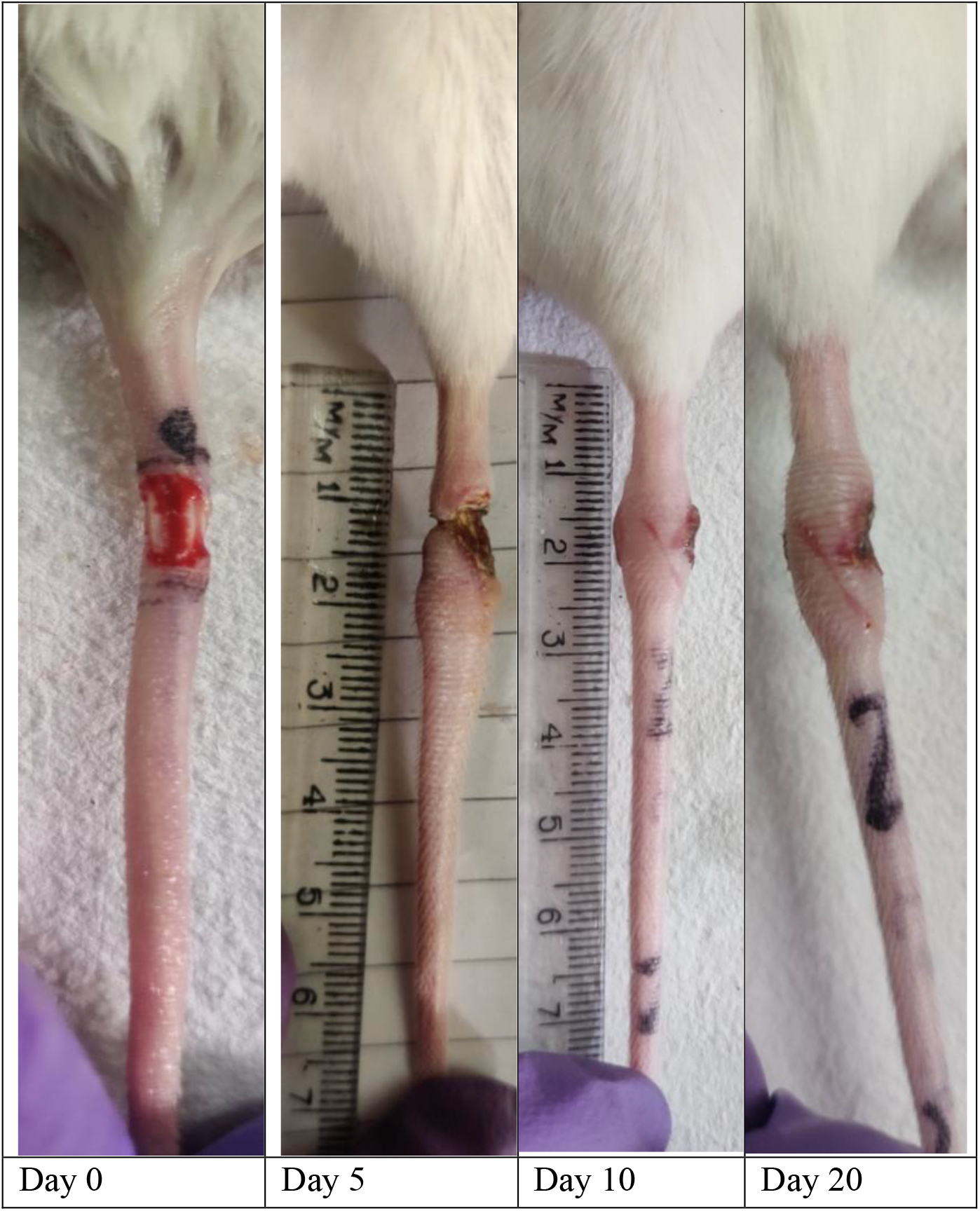
Experimental Control (n=10) group showing changes in swelling of mouse tail at different days.

**Fig.2.**
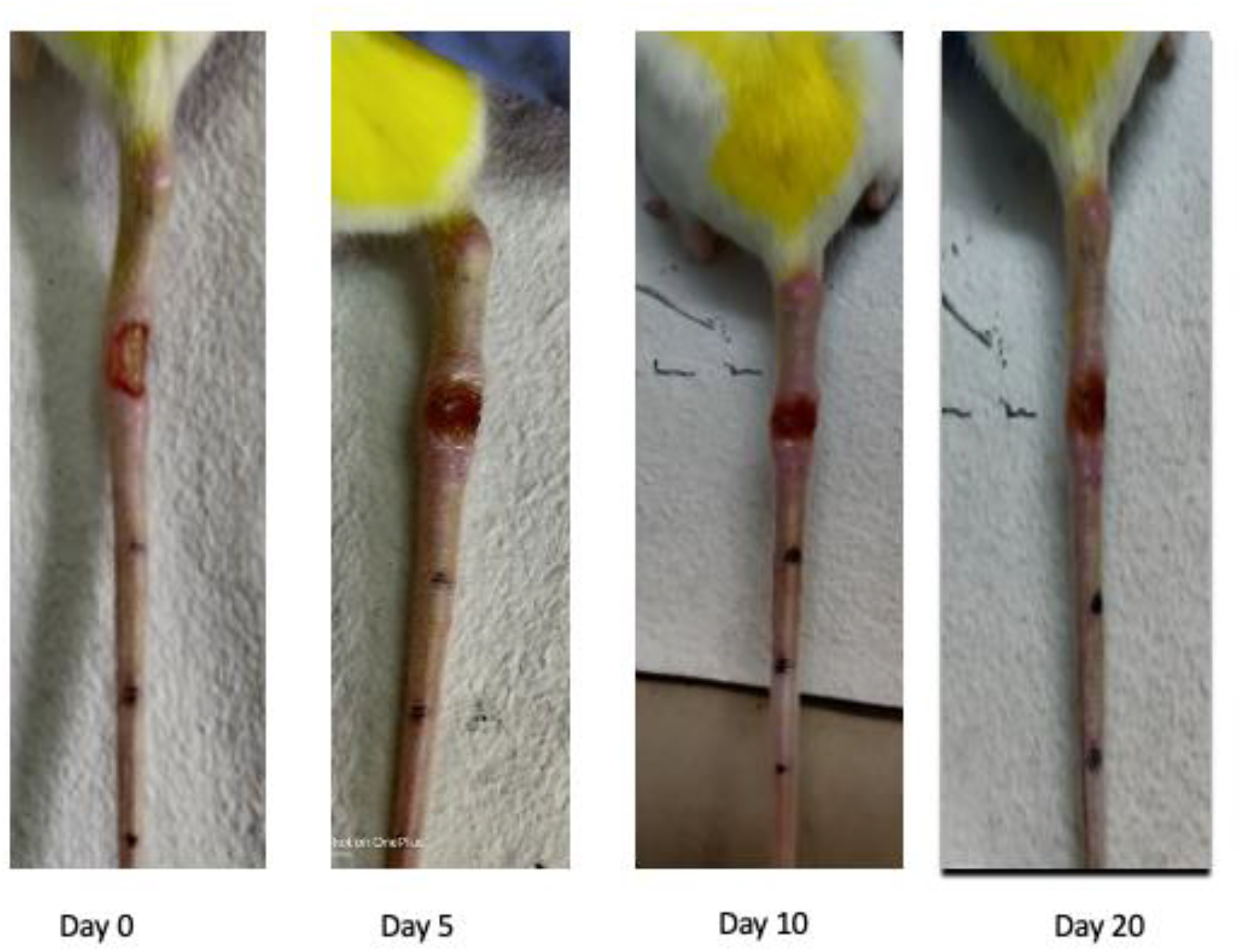
Cyperus rotundus group (n=10) showing the changes in swelling of mouse tail at different days.

**TABLE 1.**
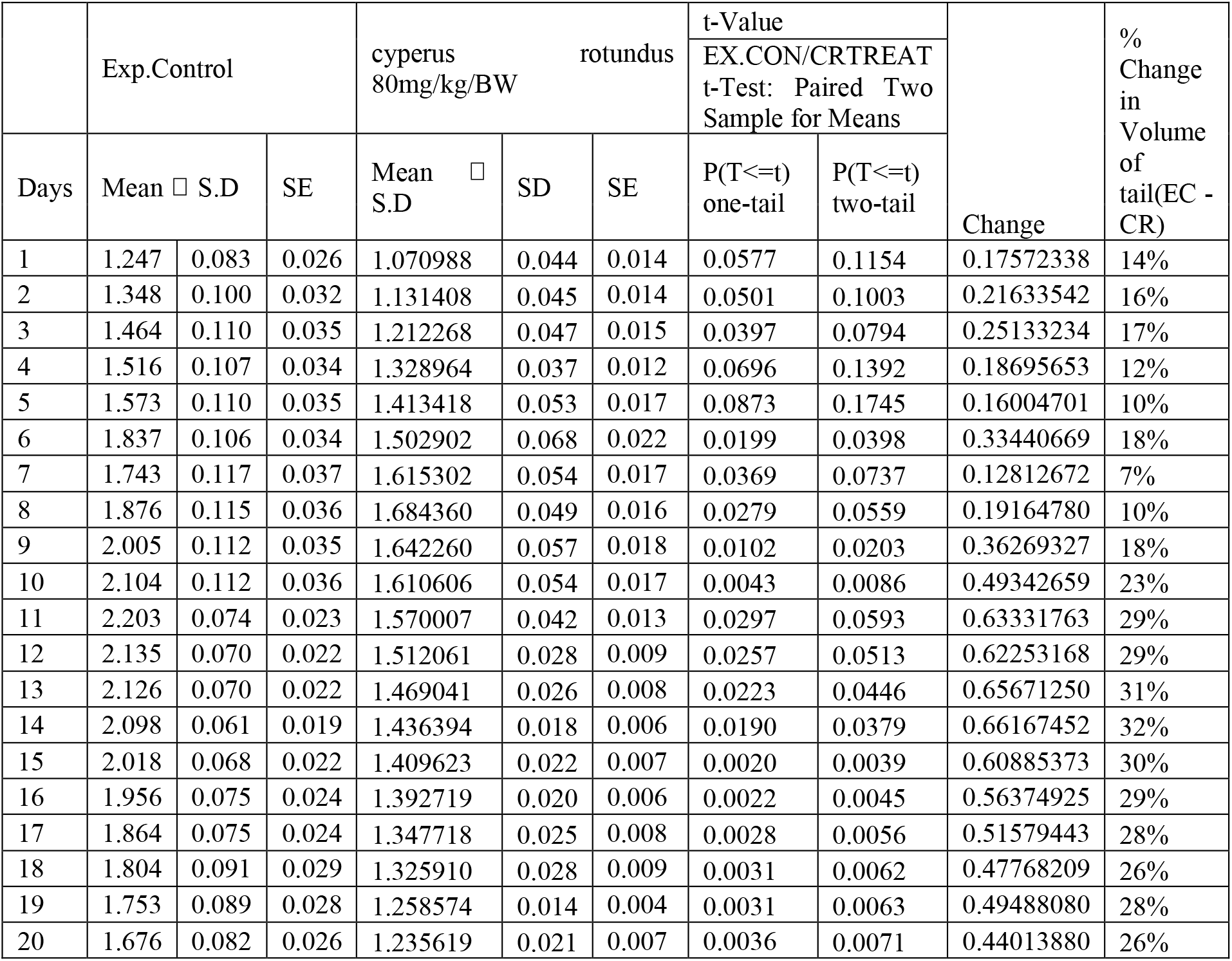
Kinetics of total volume in cut region of mice tail in Exp.Control and Cyperus rotundus.

**Fig.3.**
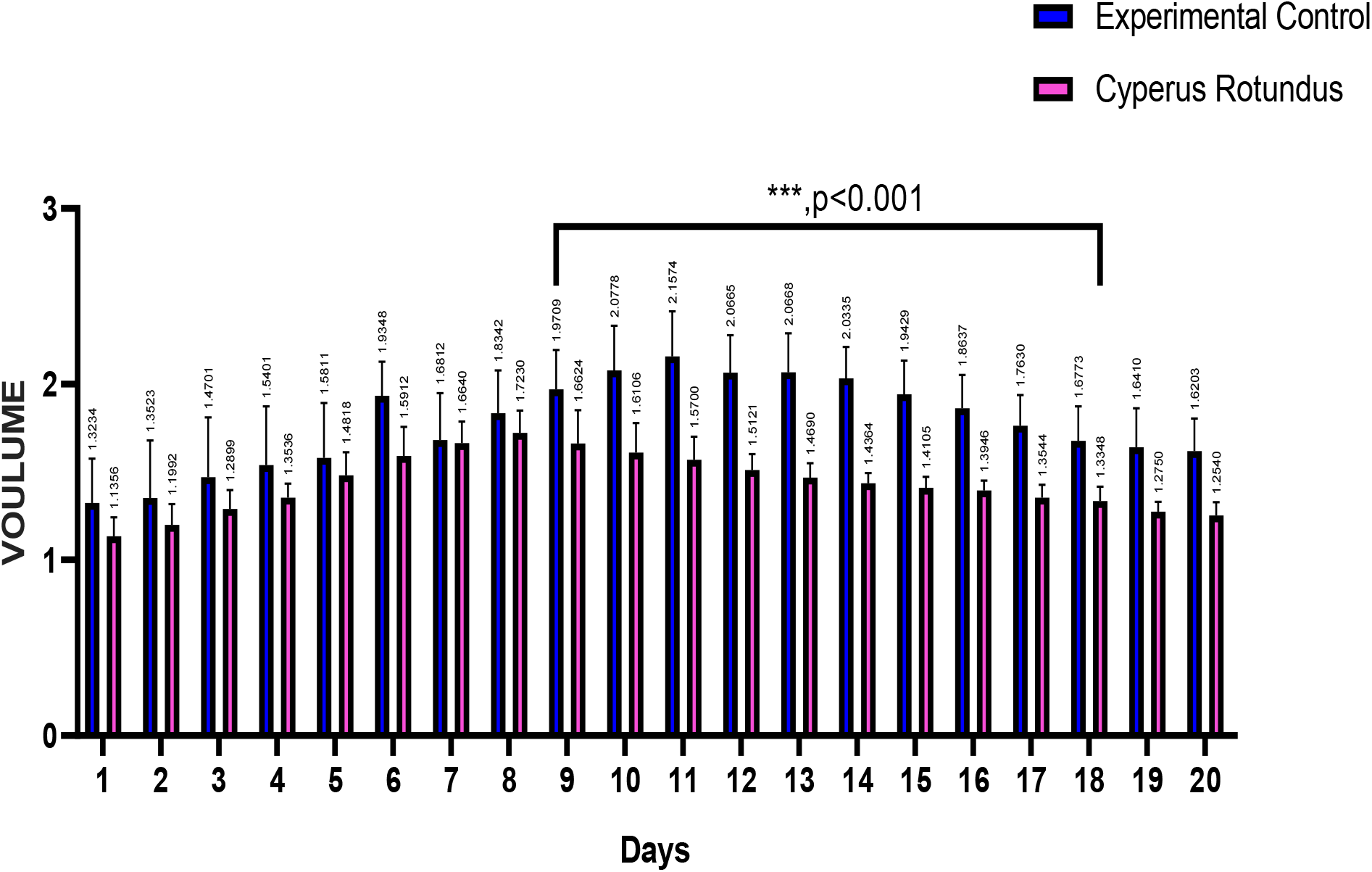
Graphical presentation of kinetical changes in the swelling of Exp.Control verses Cyperus rotundus. the Statistical analysis with Student’s t-test. The results were considered significant at *p*<0.01.

**TABLE 2.**
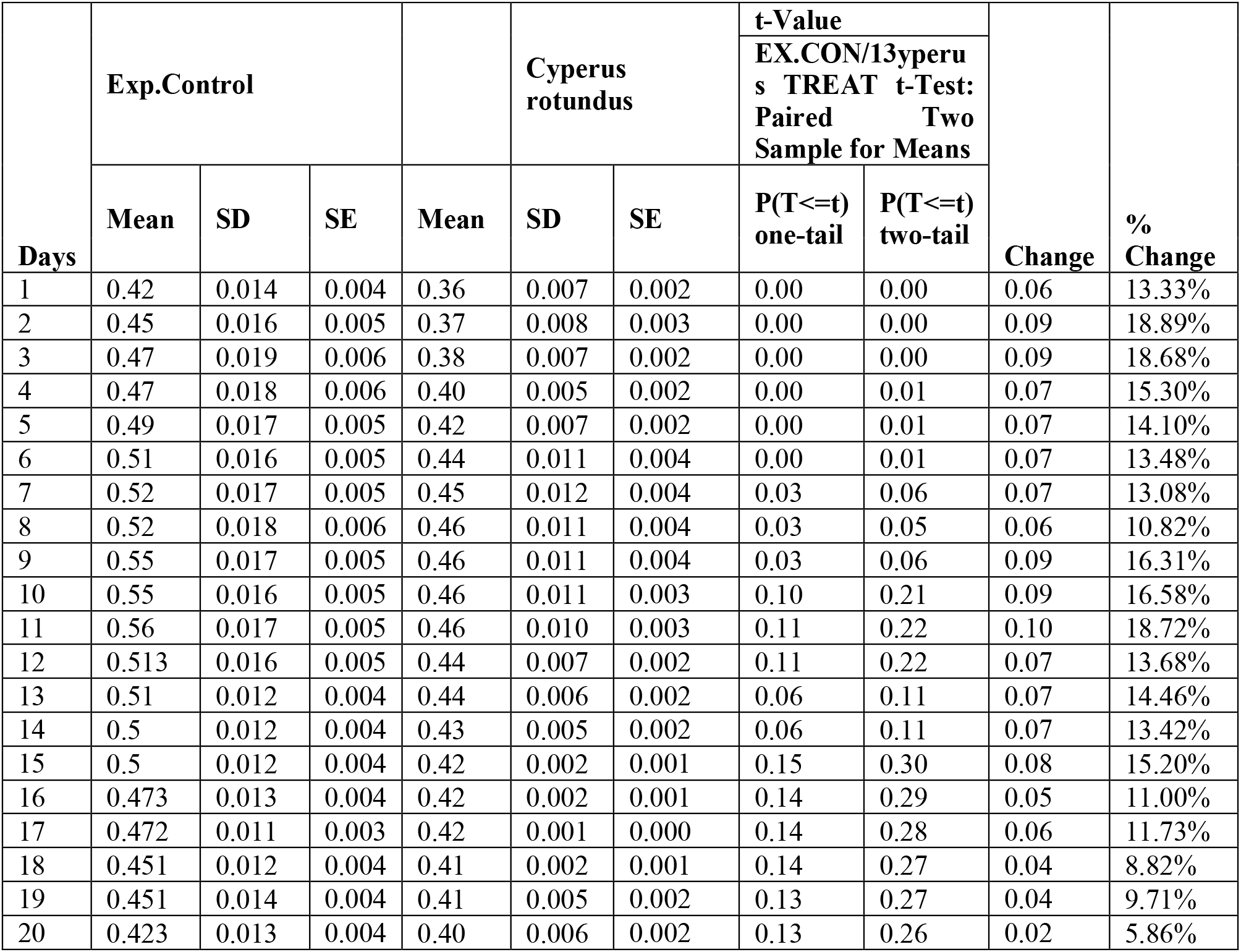
Kinetics of Upper circumference in cut region of mice tail in Exp.Control and Cyperus rotundus.

**Fig.4.**
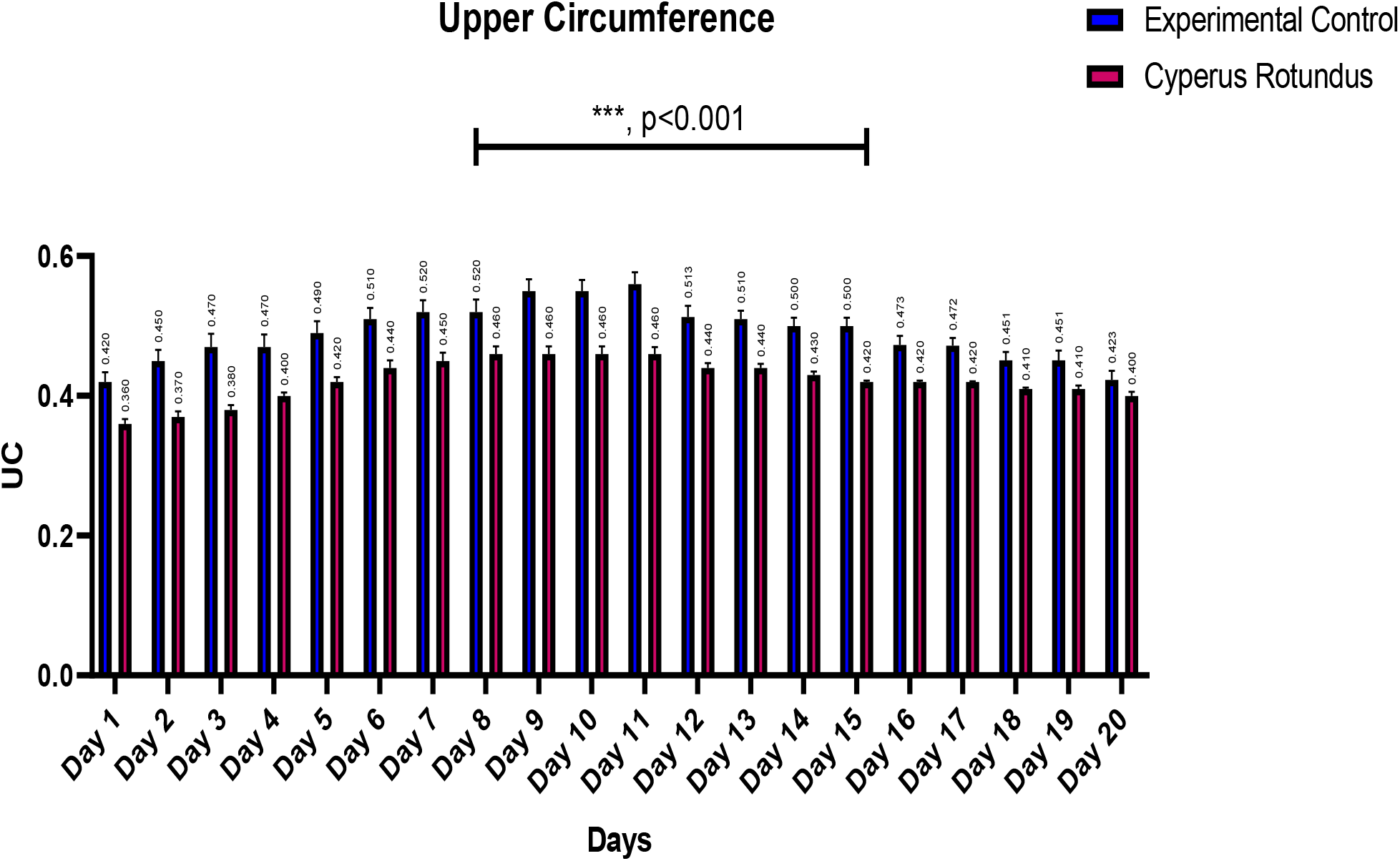
Graphical presentation of kinetical changes in the Upper circumference swelling of Exp.Control verses Cyperus rotundus. the Statistical analysis with Student’s t-test. The results were considered significant at *p*<0.001.

**TABLE 3.**
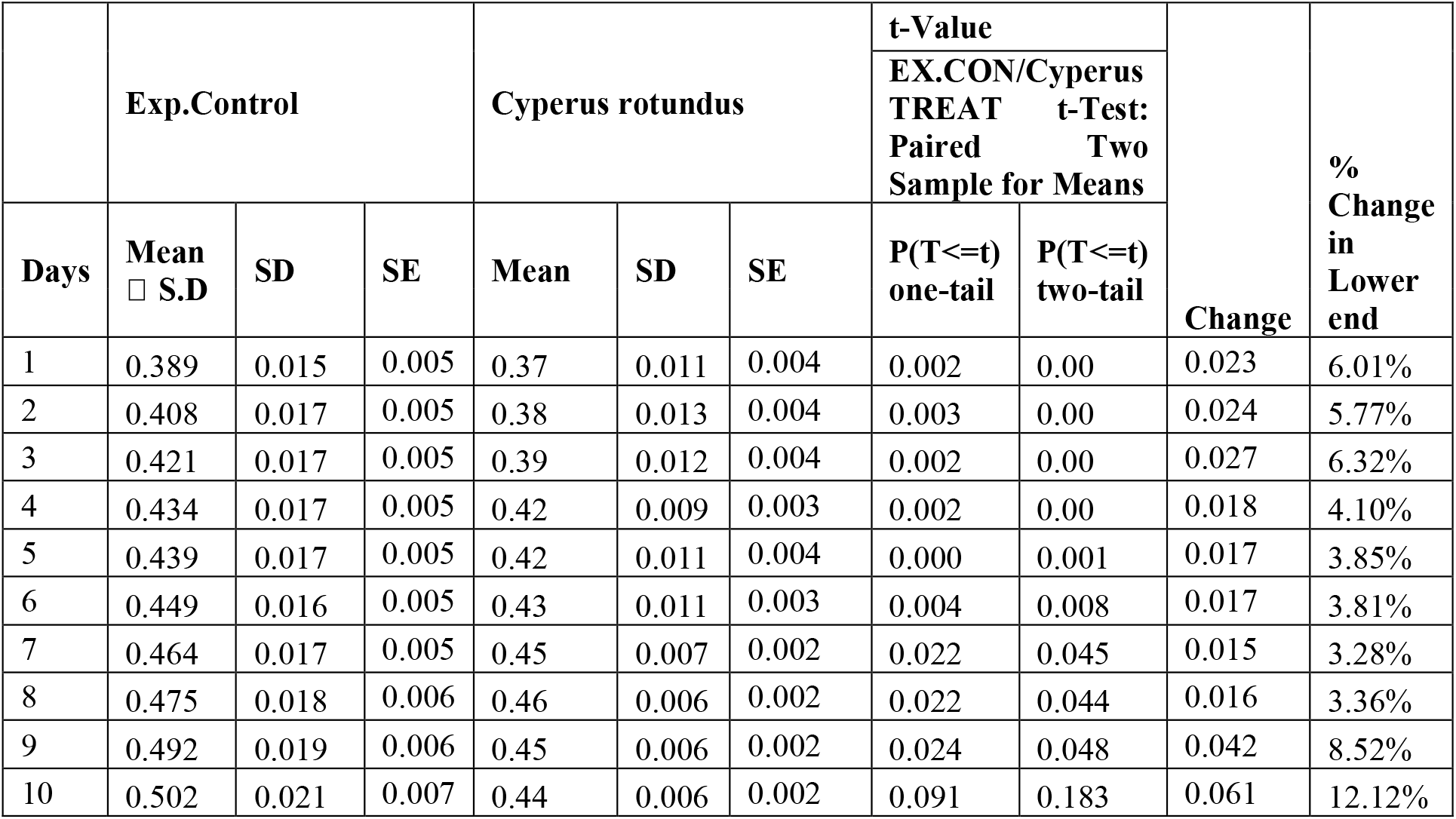
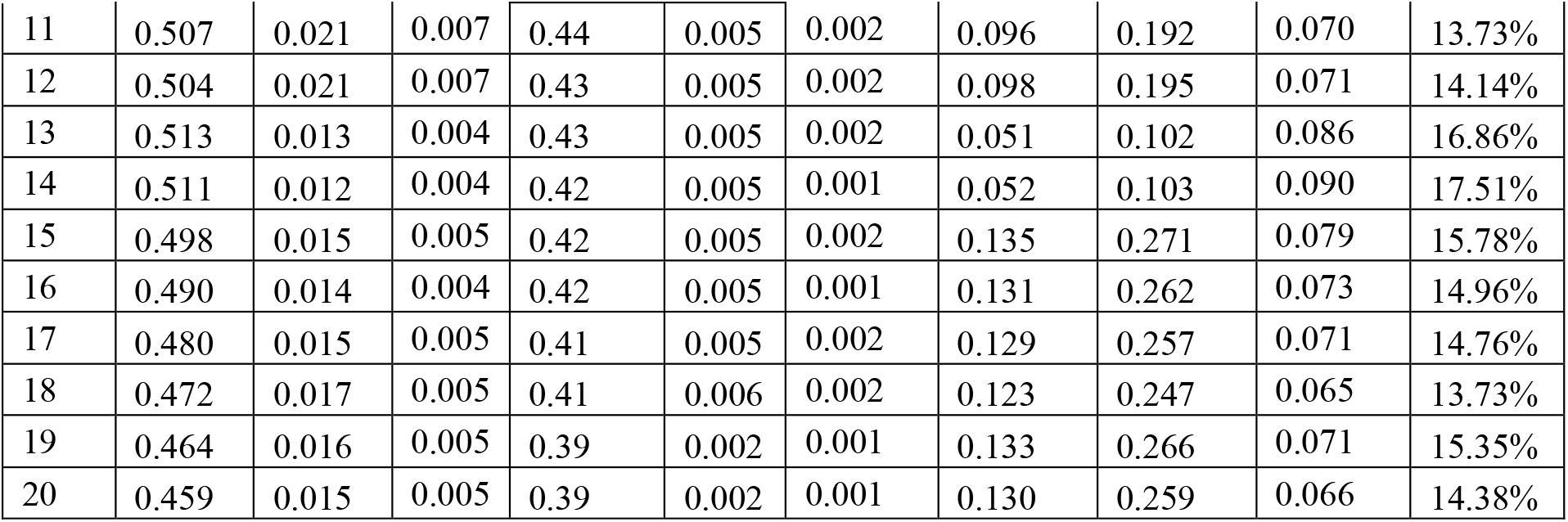
Kinetics of lower circumference in cut region of mice tail in Exp.Control and Cyperus rotundus.

**Fig.5.**
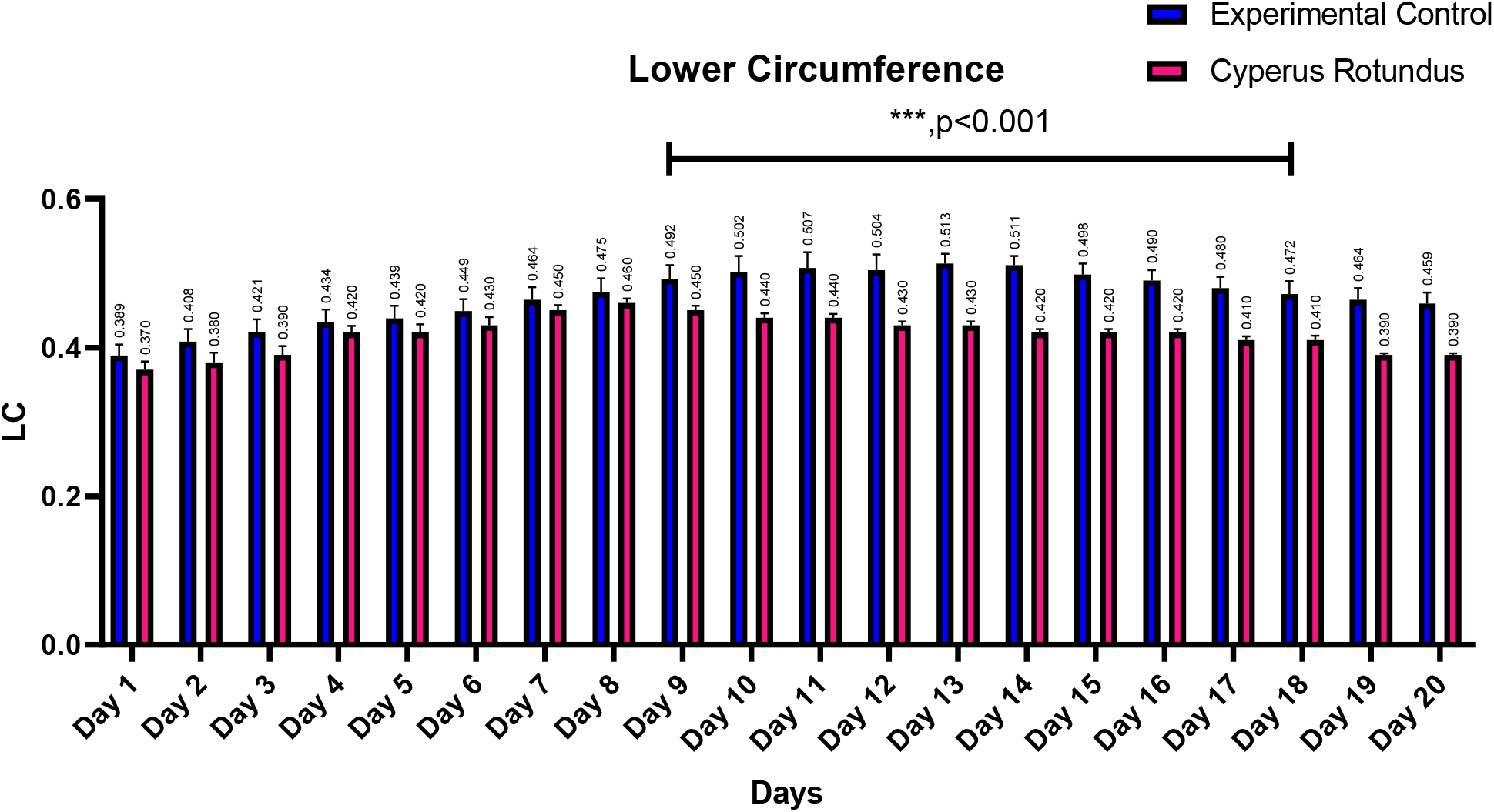
Graphical presentation of kinetical changes in the lower circumference swelling of Exp.Control verses Cyperus rotundus. the Statistical analysis with Student’s t-test. The results were considered significant at *p*<0.001.

**Table.No.4.**
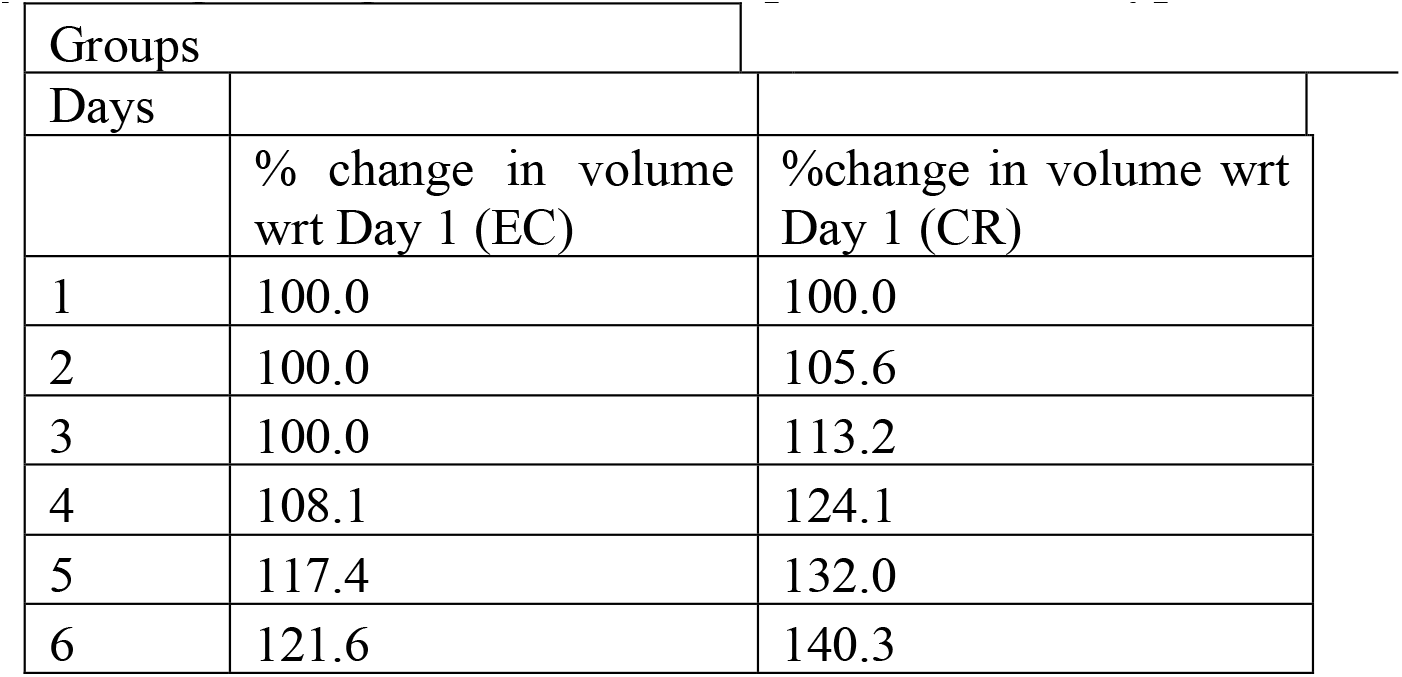
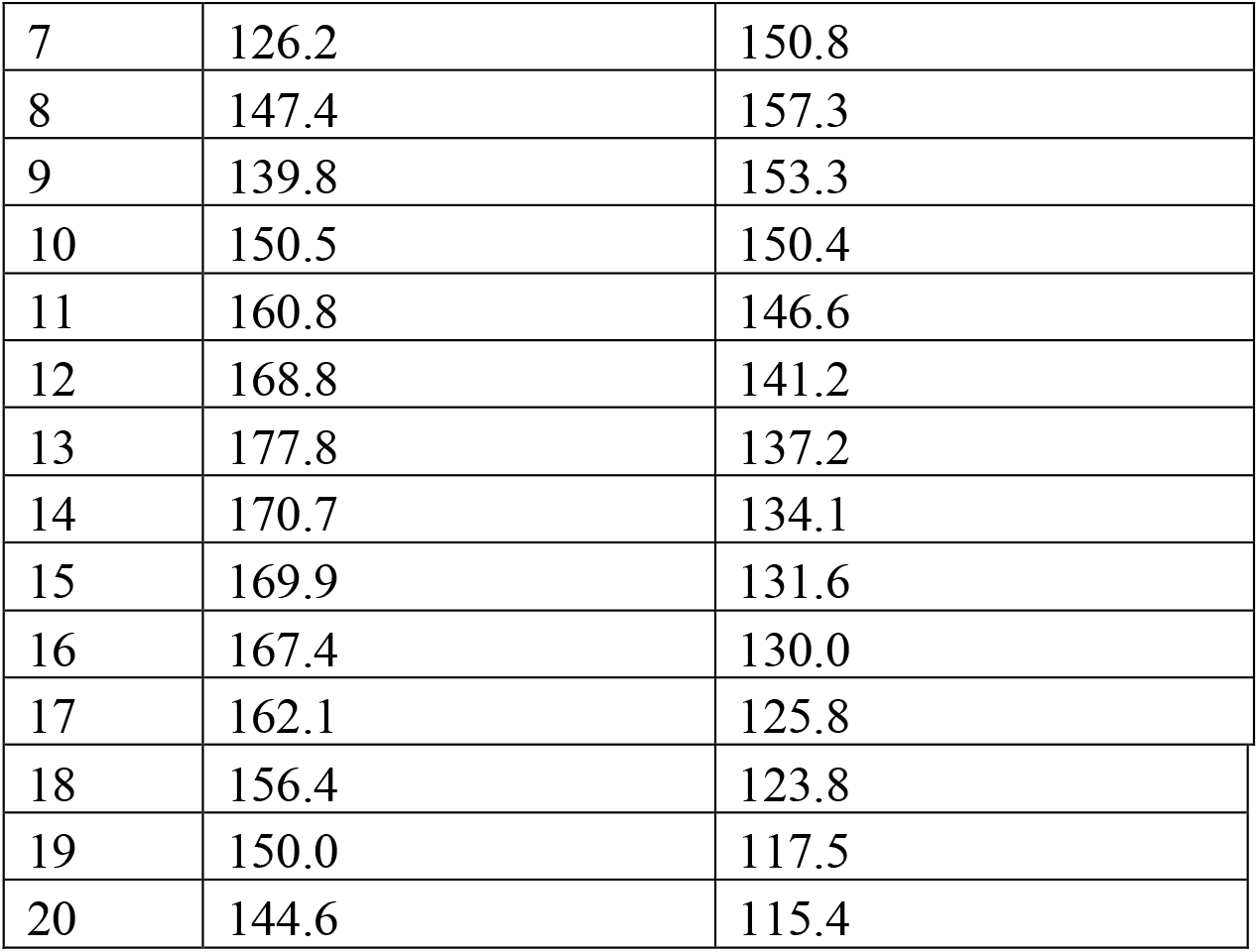
percentage change in volume in Exp.Control and Cyperus rotundus groups.

**Fig.6.**
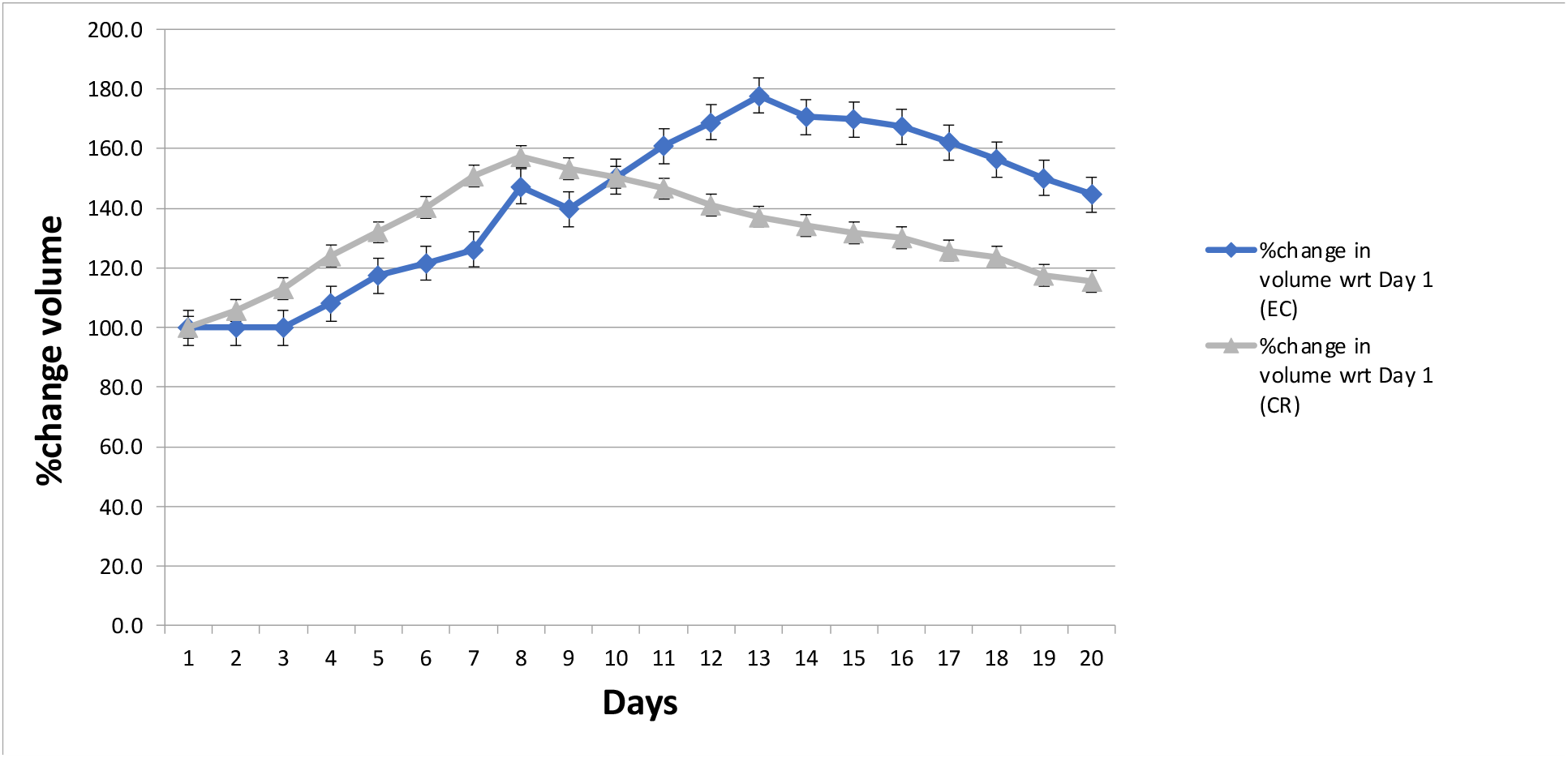
Graphical presentation of percentage change in total volume swelling of Exp.Control (blue line) and Cyperus rotundus root extract (Grey line). Each Group has been plotted w.r.t to day 1 in each of their respective group.

**Table 5.**
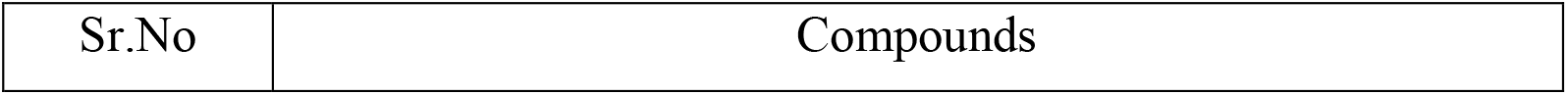
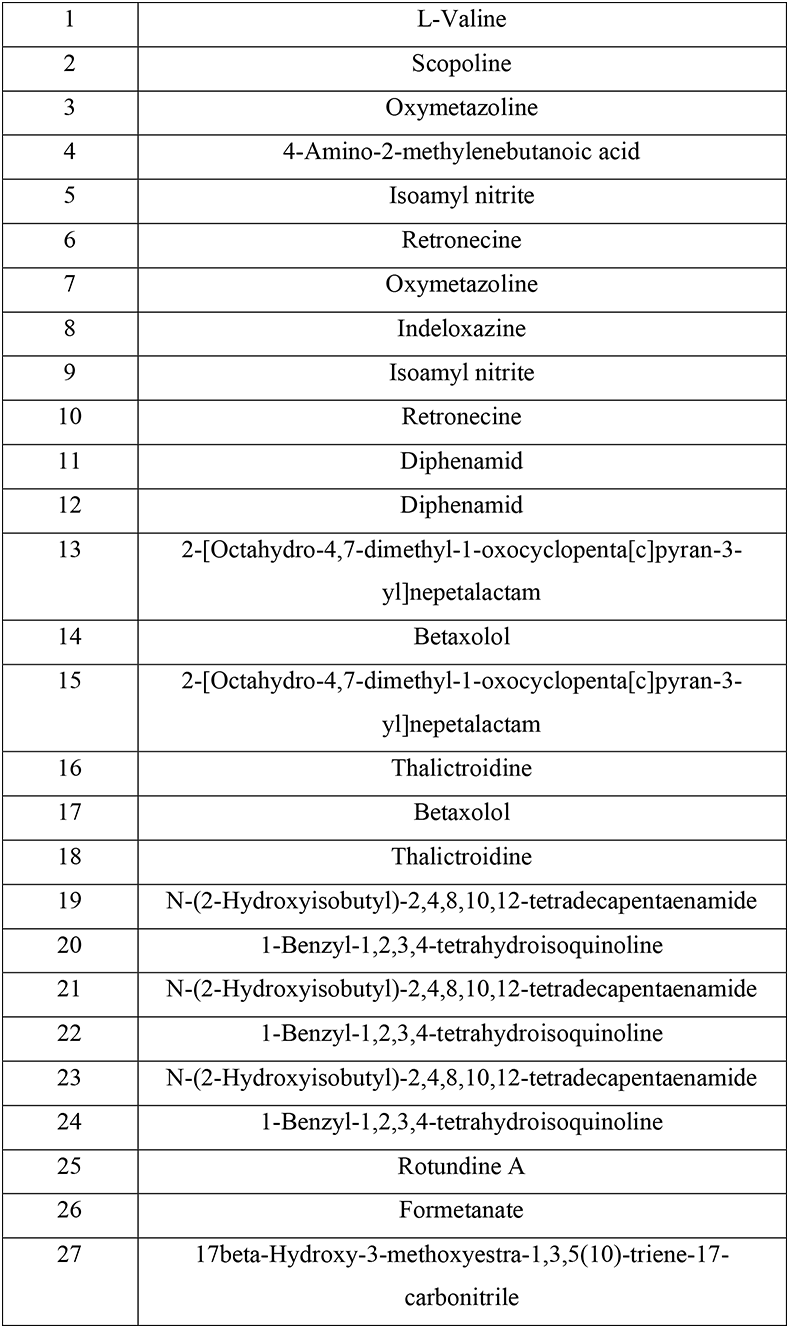
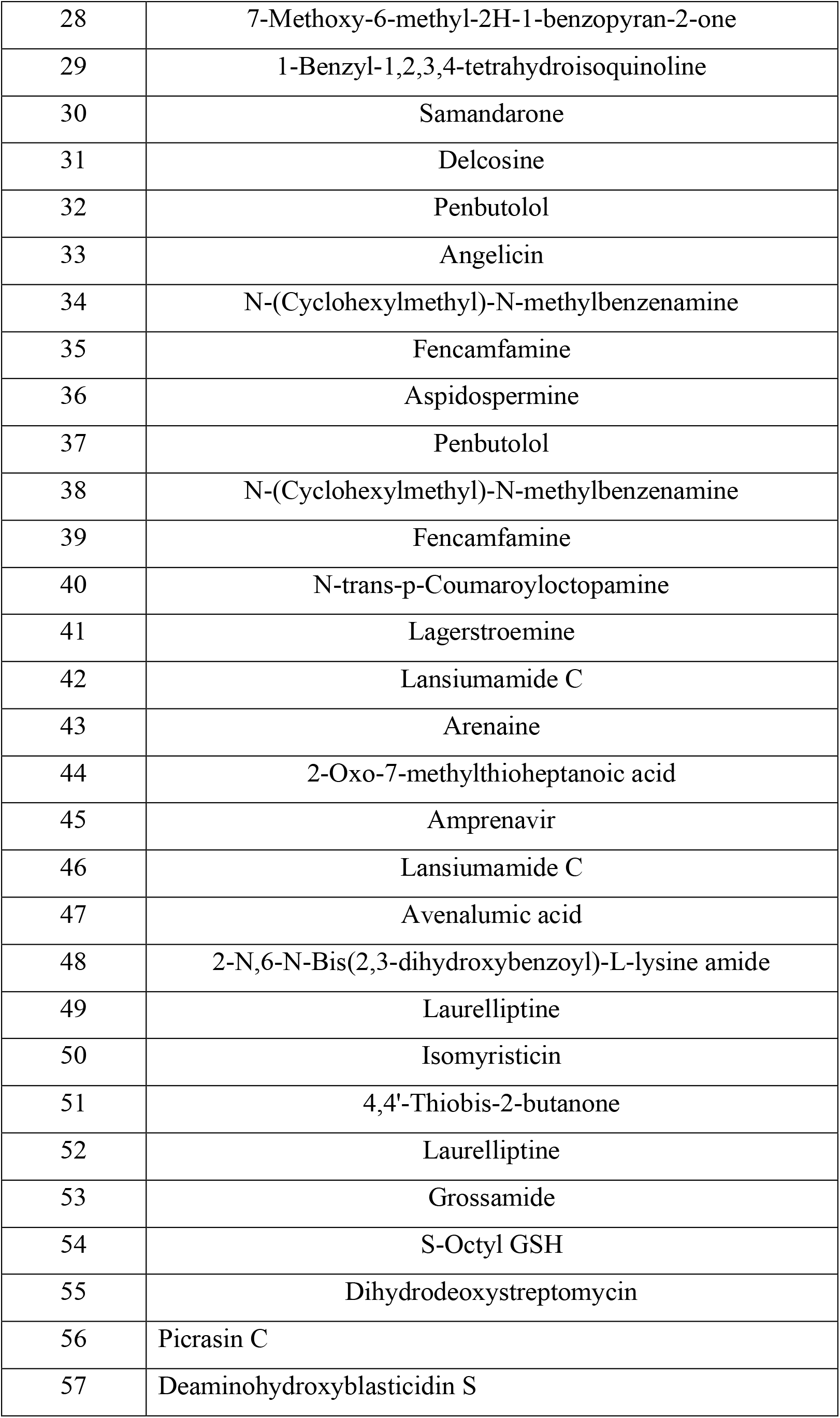
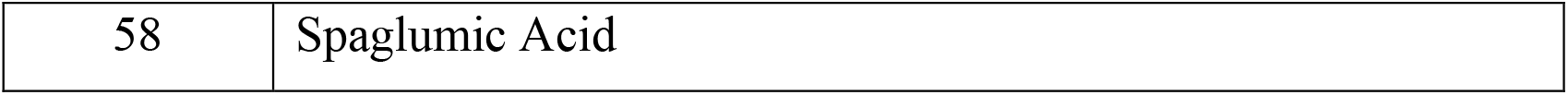
Chemical analysis of Cyperus rotundus by QTOF

**Fig.7.**
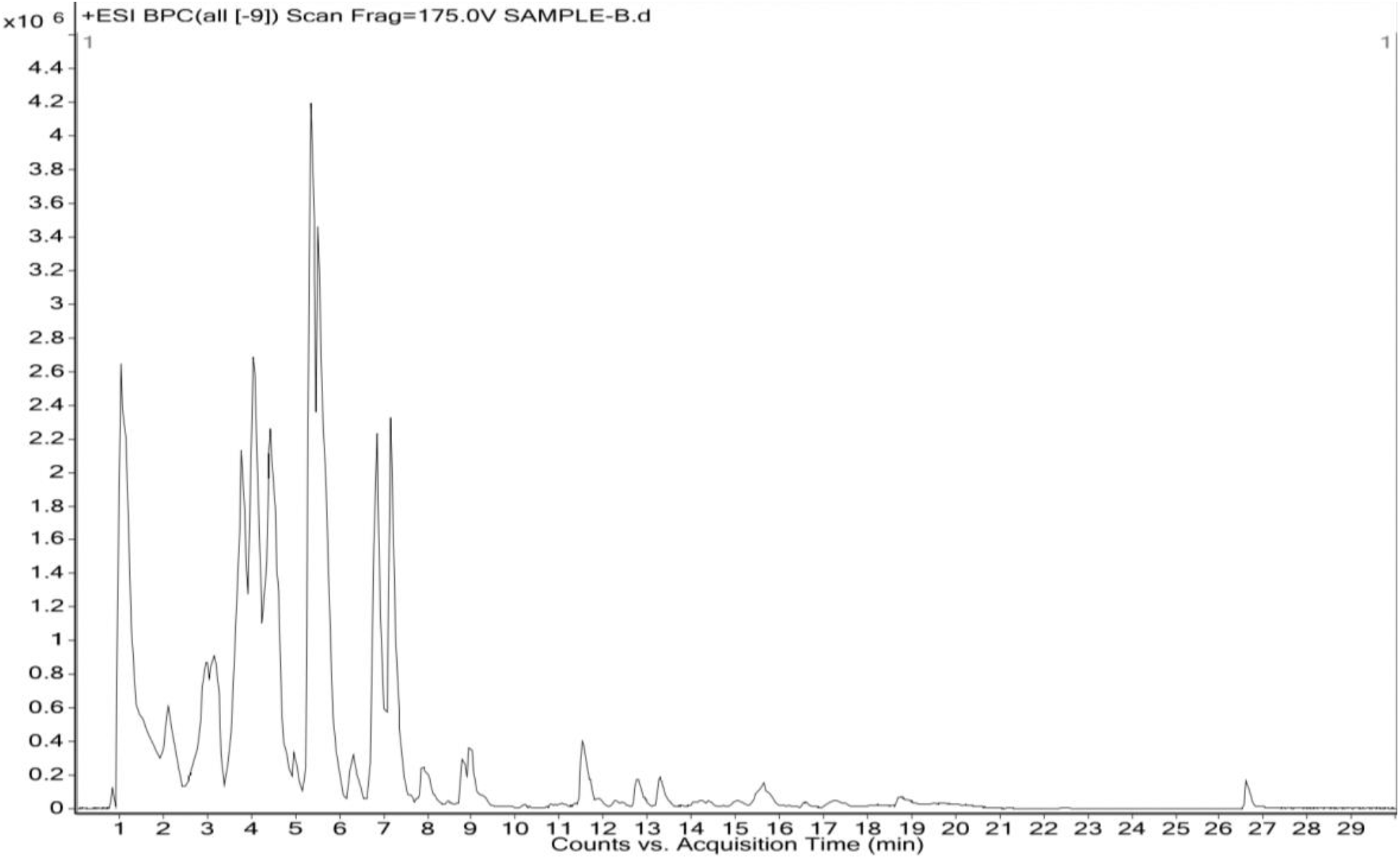
Chromatogram of QTOF(Data of Cyperus rotundus)

**Fig.8.**
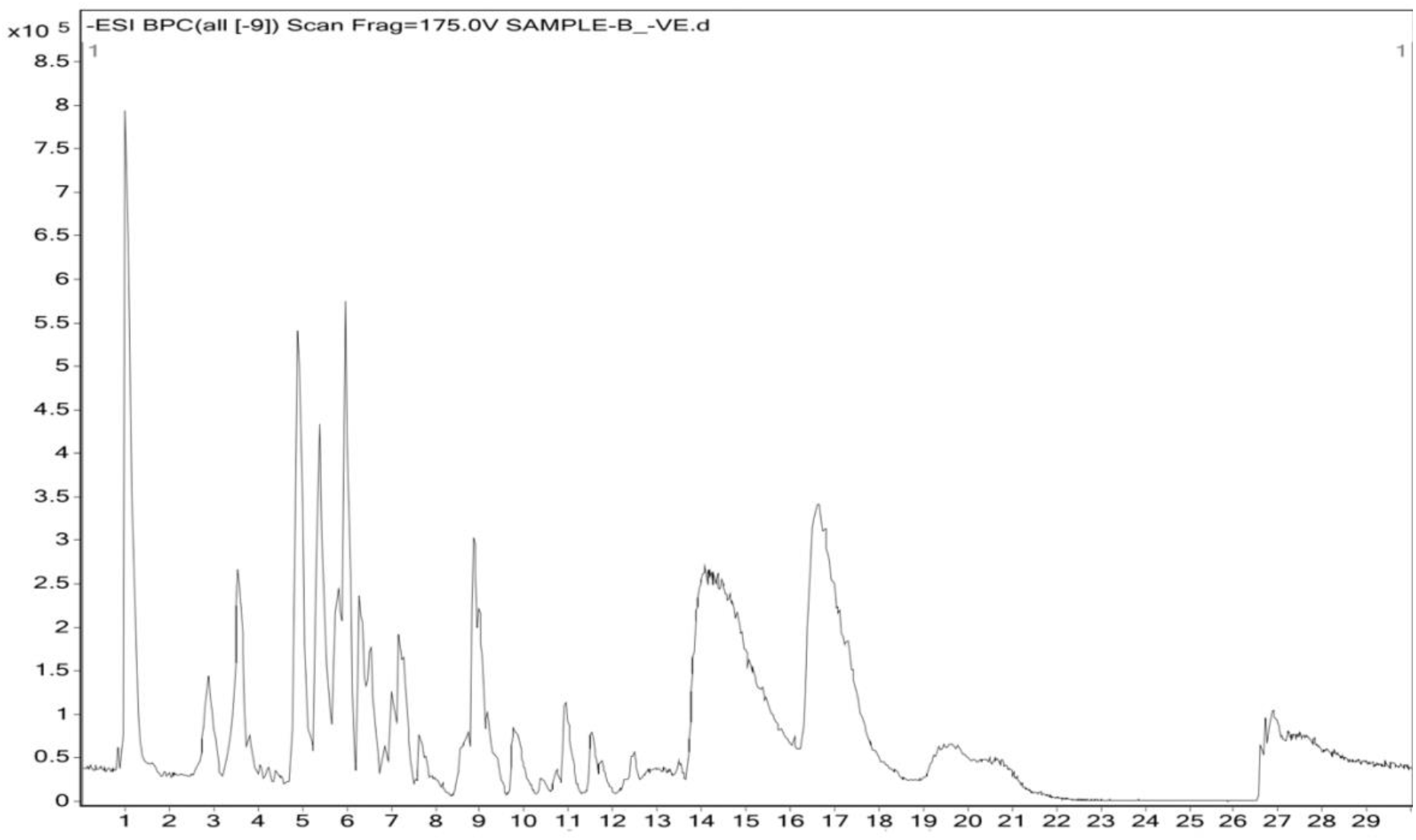
Chromatogram of QTOF(Data of Cyperus rotundus)

