## supplemental data for "Cyperus rotundus root extract inhibits progress of lymphedema in mouse tail model"

Supplementary data .

**TABLE 1. Kinetics of total volume in cut region of mice tail in Exp.Control and Cyperus rotundus**

|  | Exp.Control | | | cyperus rotundus 80mg/kg/BW | | | t-Value | | Change | % Change in Volume of tail(EC -CR) |
| --- | --- | --- | --- | --- | --- | --- | --- | --- | --- | --- |
|  |  |  |  |  |  |  | EX.CON/CRTREAT t-Test: Paired Two Sample for Means | |  |  |
| Days | Mean ± S.D | | SE | Mean ± S.D | SD | SE | P(T<=t) one-tail | P(T<=t) two-tail |  |  |
| 1 | 1.247 | 0.083 | 0.026 | 1.070988 | 0.044 | 0.014 | 0.0577 | 0.1154 | 0.17572338 | 14% |
| 2 | 1.348 | 0.100 | 0.032 | 1.131408 | 0.045 | 0.014 | 0.0501 | 0.1003 | 0.21633542 | 16% |
| 3 | 1.464 | 0.110 | 0.035 | 1.212268 | 0.047 | 0.015 | 0.0397 | 0.0794 | 0.25133234 | 17% |
| 4 | 1.516 | 0.107 | 0.034 | 1.328964 | 0.037 | 0.012 | 0.0696 | 0.1392 | 0.18695653 | 12% |
| 5 | 1.573 | 0.110 | 0.035 | 1.413418 | 0.053 | 0.017 | 0.0873 | 0.1745 | 0.16004701 | 10% |
| 6 | 1.837 | 0.106 | 0.034 | 1.502902 | 0.068 | 0.022 | 0.0199 | 0.0398 | 0.33440669 | 18% |
| 7 | 1.743 | 0.117 | 0.037 | 1.615302 | 0.054 | 0.017 | 0.0369 | 0.0737 | 0.12812672 | 7% |
| 8 | 1.876 | 0.115 | 0.036 | 1.684360 | 0.049 | 0.016 | 0.0279 | 0.0559 | 0.19164780 | 10% |
| 9 | 2.005 | 0.112 | 0.035 | 1.642260 | 0.057 | 0.018 | 0.0102 | 0.0203 | 0.36269327 | 18% |
| 10 | 2.104 | 0.112 | 0.036 | 1.610606 | 0.054 | 0.017 | 0.0043 | 0.0086 | 0.49342659 | 23% |
| 11 | 2.203 | 0.074 | 0.023 | 1.570007 | 0.042 | 0.013 | 0.0297 | 0.0593 | 0.63331763 | 29% |
| 12 | 2.135 | 0.070 | 0.022 | 1.512061 | 0.028 | 0.009 | 0.0257 | 0.0513 | 0.62253168 | 29% |
| 13 | 2.126 | 0.070 | 0.022 | 1.469041 | 0.026 | 0.008 | 0.0223 | 0.0446 | 0.65671250 | 31% |
| 14 | 2.098 | 0.061 | 0.019 | 1.436394 | 0.018 | 0.006 | 0.0190 | 0.0379 | 0.66167452 | 32% |
| 15 | 2.018 | 0.068 | 0.022 | 1.409623 | 0.022 | 0.007 | 0.0020 | 0.0039 | 0.60885373 | 30% |
| 16 | 1.956 | 0.075 | 0.024 | 1.392719 | 0.020 | 0.006 | 0.0022 | 0.0045 | 0.56374925 | 29% |
| 17 | 1.864 | 0.075 | 0.024 | 1.347718 | 0.025 | 0.008 | 0.0028 | 0.0056 | 0.51579443 | 28% |
| 18 | 1.804 | 0.091 | 0.029 | 1.325910 | 0.028 | 0.009 | 0.0031 | 0.0062 | 0.47768209 | 26% |
| 19 | 1.753 | 0.089 | 0.028 | 1.258574 | 0.014 | 0.004 | 0.0031 | 0.0063 | 0.49488080 | 28% |
| 20 | 1.676 | 0.082 | 0.026 | 1.235619 | 0.021 | 0.007 | 0.0036 | 0.0071 | 0.44013880 | 26% |

**TABLE 2. Kinetics of Upper circumference in cut region of mice tail in Exp.Control and Cyperus rotundus**

| **Days** | **Exp.Control** | | |  | **Cyperus rotundus** | | **t-Value** | | **Change** | **% Change** |
| --- | --- | --- | --- | --- | --- | --- | --- | --- | --- | --- |
|  |  |  |  |  |  |  | **EX.CON/yperus TREAT t-Test: Paired Two Sample for Means** | |  |  |
|  | **Mean** | **SD** | **SE** | **Mean** | **SD** | **SE** | **P(T<=t) one-tail** | **P(T<=t) two-tail** |  |  |
| 1 | 0.42 | 0.014 | 0.004 | 0.36 | 0.007 | 0.002 | 0.00 | 0.00 | 0.06 | 13.33% |
| 2 | 0.45 | 0.016 | 0.005 | 0.37 | 0.008 | 0.003 | 0.00 | 0.00 | 0.09 | 18.89% |
| 3 | 0.47 | 0.019 | 0.006 | 0.38 | 0.007 | 0.002 | 0.00 | 0.00 | 0.09 | 18.68% |
| 4 | 0.47 | 0.018 | 0.006 | 0.40 | 0.005 | 0.002 | 0.00 | 0.01 | 0.07 | 15.30% |
| 5 | 0.49 | 0.017 | 0.005 | 0.42 | 0.007 | 0.002 | 0.00 | 0.01 | 0.07 | 14.10% |
| 6 | 0.51 | 0.016 | 0.005 | 0.44 | 0.011 | 0.004 | 0.00 | 0.01 | 0.07 | 13.48% |
| 7 | 0.52 | 0.017 | 0.005 | 0.45 | 0.012 | 0.004 | 0.03 | 0.06 | 0.07 | 13.08% |
| 8 | 0.52 | 0.018 | 0.006 | 0.46 | 0.011 | 0.004 | 0.03 | 0.05 | 0.06 | 10.82% |
| 9 | 0.55 | 0.017 | 0.005 | 0.46 | 0.011 | 0.004 | 0.03 | 0.06 | 0.09 | 16.31% |
| 10 | 0.55 | 0.016 | 0.005 | 0.46 | 0.011 | 0.003 | 0.10 | 0.21 | 0.09 | 16.58% |
| 11 | 0.56 | 0.017 | 0.005 | 0.46 | 0.010 | 0.003 | 0.11 | 0.22 | 0.10 | 18.72% |
| 12 | 0.513 | 0.016 | 0.005 | 0.44 | 0.007 | 0.002 | 0.11 | 0.22 | 0.07 | 13.68% |
| 13 | 0.51 | 0.012 | 0.004 | 0.44 | 0.006 | 0.002 | 0.06 | 0.11 | 0.07 | 14.46% |
| 14 | 0.5 | 0.012 | 0.004 | 0.43 | 0.005 | 0.002 | 0.06 | 0.11 | 0.07 | 13.42% |
| 15 | 0.5 | 0.012 | 0.004 | 0.42 | 0.002 | 0.001 | 0.15 | 0.30 | 0.08 | 15.20% |
| 16 | 0.473 | 0.013 | 0.004 | 0.42 | 0.002 | 0.001 | 0.14 | 0.29 | 0.05 | 11.00% |
| 17 | 0.472 | 0.011 | 0.003 | 0.42 | 0.001 | 0.000 | 0.14 | 0.28 | 0.06 | 11.73% |
| 18 | 0.451 | 0.012 | 0.004 | 0.41 | 0.002 | 0.001 | 0.14 | 0.27 | 0.04 | 8.82% |
| 19 | 0.451 | 0.014 | 0.004 | 0.41 | 0.005 | 0.002 | 0.13 | 0.27 | 0.04 | 9.71% |
| 20 | 0.423 | 0.013 | 0.004 | 0.40 | 0.006 | 0.002 | 0.13 | 0.26 | 0.02 | 5.86% |

**TABLE 3. Kinetics of lower circumference in cut region of mice tail in Exp.Control and Cyperus rotundus**

|  | **Exp.Control** | | | **Cyperus rotundus** | | | **t-Value** | | **Change** | **% Change in Lower end** |
| --- | --- | --- | --- | --- | --- | --- | --- | --- | --- | --- |
|  |  |  |  |  |  |  | **EX.CON/Cyperus TREAT t-Test: Paired Two Sample for Means** | |  |  |
| **Days** | **Mean ± S.D** | **SD** | **SE** | **Mean** | **SD** | **SE** | **P(T<=t) one-tail** | **P(T<=t) two-tail** |  |  |
| 1 | 0.389 | 0.015 | 0.005 | 0.37 | 0.011 | 0.004 | 0.002 | 0.00 | 0.023 | 6.01% |
| 2 | 0.408 | 0.017 | 0.005 | 0.38 | 0.013 | 0.004 | 0.003 | 0.00 | 0.024 | 5.77% |
| 3 | 0.421 | 0.017 | 0.005 | 0.39 | 0.012 | 0.004 | 0.002 | 0.00 | 0.027 | 6.32% |
| 4 | 0.434 | 0.017 | 0.005 | 0.42 | 0.009 | 0.003 | 0.002 | 0.00 | 0.018 | 4.10% |
| 5 | 0.439 | 0.017 | 0.005 | 0.42 | 0.011 | 0.004 | 0.000 | 0.001 | 0.017 | 3.85% |
| 6 | 0.449 | 0.016 | 0.005 | 0.43 | 0.011 | 0.003 | 0.004 | 0.008 | 0.017 | 3.81% |
| 7 | 0.464 | 0.017 | 0.005 | 0.45 | 0.007 | 0.002 | 0.022 | 0.045 | 0.015 | 3.28% |
| 8 | 0.475 | 0.018 | 0.006 | 0.46 | 0.006 | 0.002 | 0.022 | 0.044 | 0.016 | 3.36% |
| 9 | 0.492 | 0.019 | 0.006 | 0.45 | 0.006 | 0.002 | 0.024 | 0.048 | 0.042 | 8.52% |
| 10 | 0.502 | 0.021 | 0.007 | 0.44 | 0.006 | 0.002 | 0.091 | 0.183 | 0.061 | 12.12% |
| 11 | 0.507 | 0.021 | 0.007 | 0.44 | 0.005 | 0.002 | 0.096 | 0.192 | 0.070 | 13.73% |
| 12 | 0.504 | 0.021 | 0.007 | 0.43 | 0.005 | 0.002 | 0.098 | 0.195 | 0.071 | 14.14% |
| 13 | 0.513 | 0.013 | 0.004 | 0.43 | 0.005 | 0.002 | 0.051 | 0.102 | 0.086 | 16.86% |
| 14 | 0.511 | 0.012 | 0.004 | 0.42 | 0.005 | 0.001 | 0.052 | 0.103 | 0.090 | 17.51% |
| 15 | 0.498 | 0.015 | 0.005 | 0.42 | 0.005 | 0.002 | 0.135 | 0.271 | 0.079 | 15.78% |
| 16 | 0.490 | 0.014 | 0.004 | 0.42 | 0.005 | 0.001 | 0.131 | 0.262 | 0.073 | 14.96% |
| 17 | 0.480 | 0.015 | 0.005 | 0.41 | 0.005 | 0.002 | 0.129 | 0.257 | 0.071 | 14.76% |
| 18 | 0.472 | 0.017 | 0.005 | 0.41 | 0.006 | 0.002 | 0.123 | 0.247 | 0.065 | 13.73% |
| 19 | 0.464 | 0.016 | 0.005 | 0.39 | 0.002 | 0.001 | 0.133 | 0.266 | 0.071 | 15.35% |
| 20 | 0.459 | 0.015 | 0.005 | 0.39 | 0.002 | 0.001 | 0.130 | 0.259 | 0.066 | 14.38% |

**Table.No.4 – percentage change in volume in Exp.Control and Cyperus rotundus groups .**

| Groups | |  | |
| --- | --- | --- | --- |
| Days |  | |  |
|  | % change in volume wrt Day 1 (EC) | | %change in volume wrt Day 1 (CR) |
| 1 | 100.0 | | 100.0 |
| 2 | 100.0 | | 105.6 |
| 3 | 100.0 | | 113.2 |
| 4 | 108.1 | | 124.1 |
| 5 | 117.4 | | 132.0 |
| 6 | 121.6 | | 140.3 |
| 7 | 126.2 | | 150.8 |
| 8 | 147.4 | | 157.3 |
| 9 | 139.8 | | 153.3 |
| 10 | 150.5 | | 150.4 |
| 11 | 160.8 | | 146.6 |
| 12 | 168.8 | | 141.2 |
| 13 | 177.8 | | 137.2 |
| 14 | 170.7 | | 134.1 |
| 15 | 169.9 | | 131.6 |
| 16 | 167.4 | | 130.0 |
| 17 | 162.1 | | 125.8 |
| 18 | 156.4 | | 123.8 |
| 19 | 150.0 | | 117.5 |
| 20 | 144.6 | | 115.4 |

Table 5.Chemical analysis of Cyperus rotundus by QTOF

| Sr.No | Compounds |
| --- | --- |
| 1 | L-Valine |
| 2 | Scopoline |
| 3 | Oxymetazoline |
| 4 | 4-Amino-2-methylenebutanoic acid |
| 5 | Isoamyl nitrite |
| 6 | Retronecine |
| 7 | Oxymetazoline |
| 8 | Indeloxazine |
| 9 | Isoamyl nitrite |
| 10 | Retronecine |
| 11 | Diphenamid |
| 12 | Diphenamid |
| 13 | 2-[Octahydro-4,7-dimethyl-1-oxocyclopenta[c]pyran-3-yl]nepetalactam |
| 14 | Betaxolol |
| 15 | 2-[Octahydro-4,7-dimethyl-1-oxocyclopenta[c]pyran-3-yl]nepetalactam |
| 16 | Thalictroidine |
| 17 | Betaxolol |
| 18 | Thalictroidine |
| 19 | N-(2-Hydroxyisobutyl)-2,4,8,10,12-tetradecapentaenamide |
| 20 | 1-Benzyl-1,2,3,4-tetrahydroisoquinoline |
| 21 | N-(2-Hydroxyisobutyl)-2,4,8,10,12-tetradecapentaenamide |
| 22 | 1-Benzyl-1,2,3,4-tetrahydroisoquinoline |
| 23 | N-(2-Hydroxyisobutyl)-2,4,8,10,12-tetradecapentaenamide |
| 24 | 1-Benzyl-1,2,3,4-tetrahydroisoquinoline |
| 25 | Rotundine A |
| 26 | Formetanate |
| 27 | 17beta-Hydroxy-3-methoxyestra-1,3,5(10)-triene-17-carbonitrile |
| 28 | 7-Methoxy-6-methyl-2H-1-benzopyran-2-one |
| 29 | 1-Benzyl-1,2,3,4-tetrahydroisoquinoline |
| 30 | Samandarone |
| 31 | Delcosine |
| 32 | Penbutolol |
| 33 | Angelicin |
| 34 | N-(Cyclohexylmethyl)-N-methylbenzenamine |
| 35 | Fencamfamine |
| 36 | Aspidospermine |
| 37 | Penbutolol |
| 38 | N-(Cyclohexylmethyl)-N-methylbenzenamine |
| 39 | Fencamfamine |
| 40 | N-trans-p-Coumaroyloctopamine |
| 41 | Lagerstroemine |
| 42 | Lansiumamide C |
| 43 | Arenaine |
| 44 | 2-Oxo-7-methylthioheptanoic acid |
| 45 | Amprenavir |
| 46 | Lansiumamide C |
| 47 | Avenalumic acid |
| 48 | 2-N,6-N-Bis(2,3-dihydroxybenzoyl)-L-lysine amide |
| 49 | Laurelliptine |
| 50 | Isomyristicin |
| 51 | 4,4'-Thiobis-2-butanone |
| 52 | Laurelliptine |
| 53 | Grossamide |
| 54 | S-Octyl GSH |
| 55 | Dihydrodeoxystreptomycin |
| 56 | Picrasin C |
| 57 | Deaminohydroxyblasticidin S |
| 58 | Spaglumic Acid |
